# Noise-driven morphogenesis independent of transcriptional regulatory programs

**DOI:** 10.1101/2025.08.04.662975

**Authors:** Jack Toppen, Saeed Tavazoie

## Abstract

Development is widely understood as a deterministic process driven by transcriptional programs that specify cell fate and orchestrate morphogenesis. However, this view overlooks pervasive stochasticity in gene expression, often considered an obstacle to reliable tissue patterning. Here, we introduce stochastic tuning-driven morphogenesis (STM), an alternative conceptualization of development in which noisy gene expression is not a nuisance but the primary driving force—guiding cell fates toward optimal multicellular configurations by a trial-and-error process analogous to reinforcement learning. STM operates independently of fixed transcriptional programs, instead relying on convergence of sensory information into signaling hubs, which by reinforcing random transcriptional changes, prospectively and contextually fine-tune gene expression along key developmental milestones. STM offers a fundamentally different view of development—one in which stochastic gene expression enables real-time optimization of gene expression toward multicellular objectives, implementing a self-organizing process that is inherently resistant to molecular and environmental fluctuations.

## Introduction

Despite more than a century of developmental biology, many aspects of metazoan morphogenesis remain profoundly mysterious. There is a vast literature on phenomenological observations in key model organisms^1^. However, we still lack a predictive understanding of pattern formation in all but a few spatiotemporally restricted systems such as the early *Drosophila* embryo^2^. Foundational work in *Drosophila* and other model systems has led to the gene regulatory network (GRN) paradigm of development^3^. In the GRN conception, developmental patterns arise from a deterministically orchestrated sequence of transcriptional regulatory events in response to local morphogen gradients, neighboring cell ligands, and mechanical forces. A fundamental principle at the core of this dominant view is that the detailed trajectories of gene expression in individual cells/nuclei are explicitly encoded at the genetic level through the organization of DNA-regulatory regions composed of transcription factor (TF) binding sites^3–5^. As such, developmental trajectories of individual cells are precisely orchestrated by a step-by-step process of sensory perception, signal-transduction, and transcriptional regulation that culminates in robust and reproducible morphogenesis.

A key feature of the GRN framework is the requirement that detailed spatiotemporal trajectories of cell states and movements are explicitly encoded by the genome. We currently lack the theoretical basis to quantify the genomic information relevant to morphogenesis. However, the deterministic nature of such a precise step-by-step encoding would impose heavy demands on the information capacity of genomes of limited size. Highly reproducible developmental outcomes clearly support precise deterministic regulatory programs. However, it is becoming increasingly clear that the remarkable robustness of development occurs despite high stochasticity at the molecular level^6–22^. This is especially true in the process of mRNA transcription which is often characterized by highly non-uniform periods of transcriptional bursting^23–25^. Is this widespread stochasticity an irreducible nuisance of molecular processes operating with low number of components, or is noise actively utilized as a driving force during development? Stochasticity is exploited to generate diversity in later stages of organ maturation, for example in establishing photoreceptor^26^ and olfactory receptor^27^ choice. However, could noise fulfill a more central role throughout all stages of development?

The question of whether all cellular behaviors are explicitly encoded by the gnome has been recently addressed by work on the adaptation of budding yeast cells to unfamiliar environments. Freddolino *et al*. have found evidence for a proposed mode of cellular adaptation, termed stochastic tuning, in which adaptive gene expression states are achieved by active reinforcement of random transcriptional changes that improve the overall health of the cell^28^ (**Fig 1**). As such, stochastic tuning, is an objective-driven process operating to maximize the health of the cell in the absence of a genetically encoded transcriptional regulatory program. A global health integrator is postulated to integrate a variety of parameters relevant to cell health and to actively reinforce random transcriptional changes that improve the overall health of the cell (**Fig 1**). Stochastic tuning, thus, enables cells to establish arbitrary gene expression states that are adaptive under unfamiliar conditions not encountered in their evolutionary history^28^. A key benefit of stochastic tuning is that the genome is not required to encode regulatory programs for an endless array of possible environmental conditions. Rather, individual cells have agency to ‘discover’ adaptive gene expression states through a health-reinforced stochastic search. Recent studies have provided evidence for stochastic tuning in fission yeast^29^ and in cancer chemotherapy adaptation^30–32^. Thus, stochastic tuning may be a widespread mechanism by which eukaryotic cells adapt to novel environments beyond the capacity of their hard-coded gene regulatory networks.

**Fig. 1.**
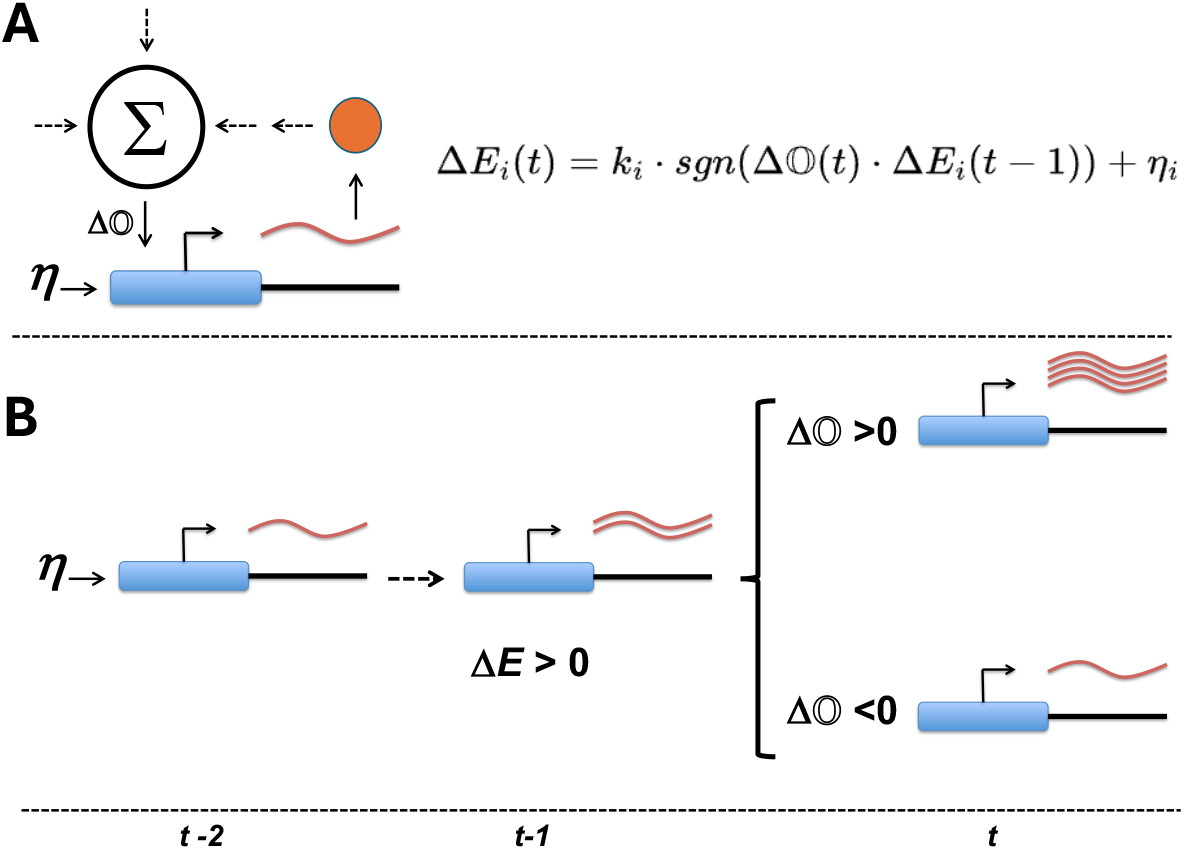
Stochastic tuning utilizes transcriptional noise and feedback from a global objective (eg. cell health) to empirically search for, and establish, optimal gene expression states. (**A**) Stochastic tuning model. Individual genes can exhibit random bursts of transcription (η). The gene maintains a temporary record (memory) of a burst through modification of local chromatin. A global health integrator (Σ) continuously broadcasts a signal (ΔO) to all promoters, conveying whether the overall objective (e.g. cell health) is improving or deteriorating. This corresponds to a form of gradient ascent optimization, where at any point, the expression-apparatus executes a change in transcriptional activity of the *i*th gene, Δ*E*_*i*_(*t*), proportional (*k*_*i*_) to the product of previous change in expression, Δ*E*_*i*_(*t-1*), and the current change in the objective, ΔO(*t*), plus discrete random noise with amplitude (*η*_*i*_). The *sgn*() function turns this into discrete effects (−1,0,1). (**B**) A simple example of stochastic tuning can be seen for a gene that experiences a random burst in transcription. If this is followed by an improvement in the overall objective (ΔO>0), the expression apparatus further increases transcriptional activity. Conversely, if there is a deterioration in the objective (ΔO<0), the expression apparatus decreases transcriptional activity. This process leads to establishment of expression levels that maximize the overall objective (e.g. health). Adapted from (Freddolino et al. *eLife* (2018) doi: 10.7554/eLife.31867).

Evidence for a noise-driven adaptation process in yeast and mammals inspired us to consider whether a multicellular version may have evolved to orchestrate gene expression trajectories during development. We wondered whether the framework of stochastic tuning could form the foundation for an alternative mode of pattern formation in which cell states are driven by multicellular objectives rather than precisely encoded step-by-step transcriptional regulatory programs (**Fig 2**). In this alternative conception, we call **S**tochastic **T**uning-driven **M**orphogenesis (STM), individual cells utilize gene expression noise and reinforcement by a multicellular objective to search for, and establish, gene expression states that increase the objective. In contrast to yeast where the sole objective is a cell’s overall health, these multicellular objectives can be thought of as ‘functions’ that integrate relevant features of the cell’s state, together with incoming sensory information from neighboring cells, to indicate optimality of multicellular configurations. The molecular realizations of these multicellular objectives correspond to signaling integration hubs that compute an overall objective whose scalar value is optimal when the cell achieves the proper gene expression state in the context of its developmentally appropriate cellular neighborhood.

**Fig. 2.**
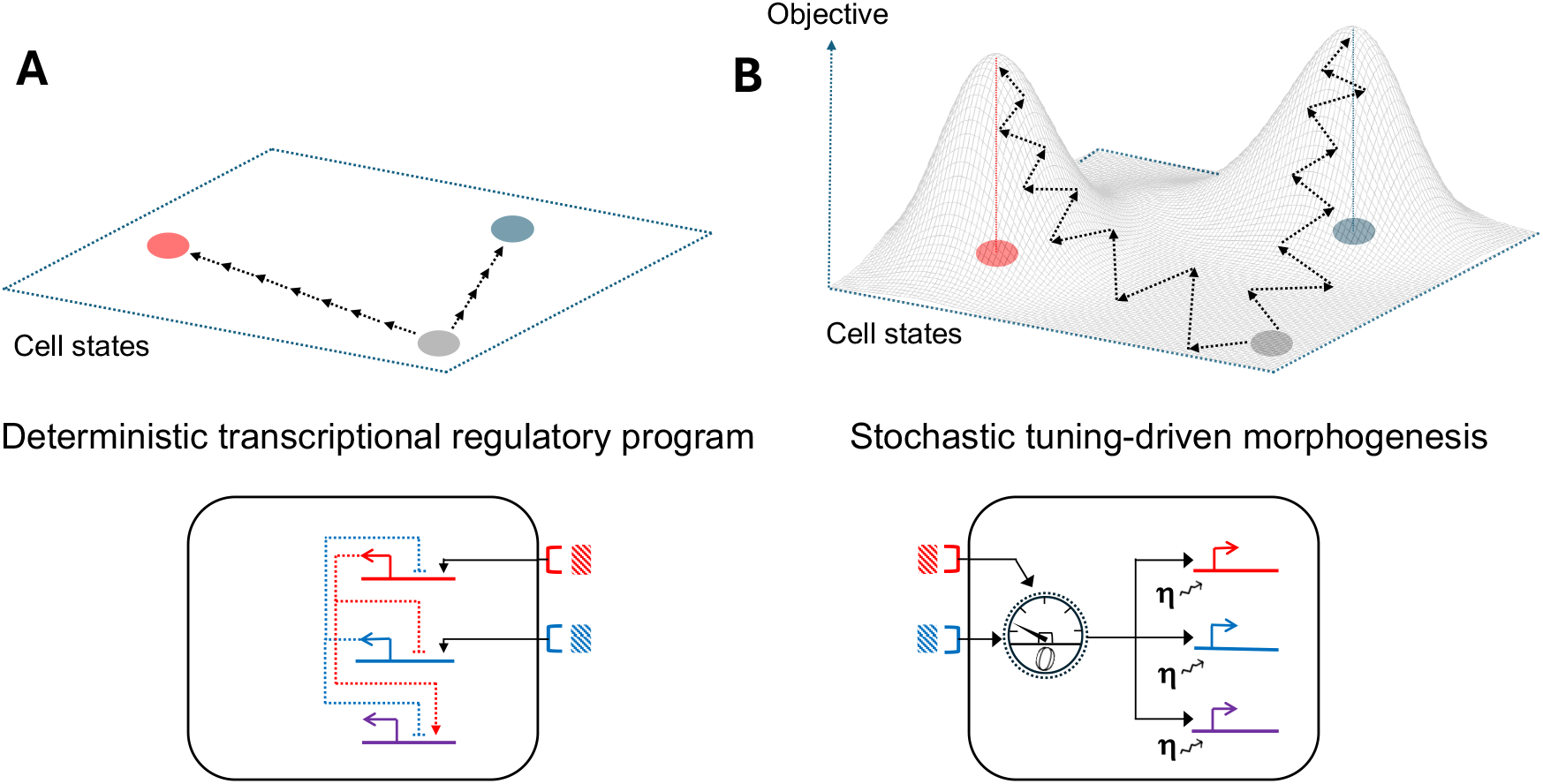
Stochastic tuning-driven morphogenesis vs. conventional transcriptional regulatory programs. (**A**) Conventional transcriptional regulatory programs execute deterministic trajectories of cell-states via activation of hard-coded regulatory circuits shown below as a network of transcriptional regulatory interactions under the influence of signaling from two external ligands. An abstract cell-state landscape depicts differentiation towards two distinct cell-states (red and blue cell types) proceeding via deterministic cascades depicted by sequential linear arrows. (**B**) In stochastic tuning-driven morphogenesis, cells utilize gene expression noise to make random changes to gene expression and reinforce those changes that increase the scalar value of a multicellular objective (***O***). The multicellular objective integrates external signaling events and internal state into computations that reflect optimal multicellular configurations. The multicellular objective has multiple optima and depending on the proximity of individual cells to these optima, and random excursions, cells can ‘flow into’ either attractor basin (red and blue cell types). Despite eventual convergence to these optima, trajectories can exhibit significant non-deterministic behavior.

In the STM framework, the genetic encoding of development does not encompass the detailed high-dimensional trajectories of every cell’s gene expression (**Fig 2A**). Rather, the genetic encoding is limited to a set of intermediate objectives reflecting key milestone multicellular configurations, and much of the transcriptional and cell behavioral dynamics between these milestones are not explicitly encoded—rather they are executed by an objective-driven search for optimality (**Fig 2B**). The STM framework can, thus, provide significant economy in the genetic encoding of morphogenesis, delegating a substantial component to a search process. Another benefit of STM is noise-resilience. Since noise is the key driving force for STM, robustness to both molecular and environmental noise is a built-in feature. Here we present the theoretical framework for STM and use simulations to demonstrate successful pattern formation across a variety of prototypic scenarios.

## Results

### A conceptual framework for stochastic tuning-driven morphogenesis

Here, we consider a toy model of pattern formation in two dimensions in which cells do not undergo cell divisions or movement, and, as such, the system can be considered an on-lattice agent-based simulation. The layer of cells is arrayed in space with coordinates (x,y), with each cell encoding a set of genes corresponding to either diffusible ligands, membrane-bound ligands, or membrane-bound receptors. The system may also be under the influence of external ligands with desired concentration profiles across the tissue (i.e. external morphogen gradients). Interactions between ligands and receptors generate signaling events that are component inputs into a signaling integration hub that, in effect, implements the multicellular objective function—henceforth referred to as the multicellular objective, or objective. As we show below, the mathematical form of these multicellular objectives maps directly to the topologies of typical signaling pathways, with crosstalk and convergence to signaling integration hubs^33–43^. In effect, the topology of these signaling pathways implement computations that reflect multi-peaked objective functions whose scalar output drives cell-states towards individual optima by stochastic tuning (**Fig 2B**).

Along with input from extracellular signaling pathways, multicellular objectives may also depend on levels of specific intracellular proteins. As such, the objective has access to the intracellular state and the sensory input converging on the cell’s surface. At any time, the objective broadcasts a single scalar output, reflecting the change in the objective relative to the previous epoch. The output of the objective, thus, reflects the specific way in which the combination of incoming signals and intracellular factors combine to generate a scalar output reflecting whether the cell is approaching a multicellular optimum, as determined by its own state and the state of the signaling events impinging on it.

Each gene has a minimal promoter, enabling its basal expression. However, there exist no conventional transcriptional regulatory input into any gene and, thus, the expression of each gene is entirely dependent on stochastic tuning (**Fig 1**) driven by transcriptional noise and reinforcement from a global multicellular objective (**Fig 2B**). If a random transcriptional change is followed by a subsequent increase in the objective, then it is reinforced. Conversely, if a random transcriptional change is followed by a decrease in the objective, then it is penalized. This simple dynamic corresponds to a form of gradient ascent^44^ that drives the expression of all the genes to levels that maximize the objective. We can equally represent optimization paths on an energy landscape where optimal low-energy configurations are achieved by gradient decent^44^. In the case of a single-cell health objective, we have shown that this mechanism can simultaneously establish arbitrary optimal gene expression states for thousands of genes^28^.

### Multicellular objectives drive cell fates toward optimal multicellular configurations

How do we construct multicellular objectives that drive desired developmental patterns? The molecular instantiation of these objectives would have emerged through evolutionary processes that accompany increasing morphological complexity of metazoans. In the absence of this historical knowledge, our challenge is to rationally engineer objectives that lead to developmental patterns consistent with what we see in nature. Our goal is, thus, to see whether such engineered objectives have the capacity, in principle, to generate complex patterns, rather than converging onto the actual naturally evolved objectives. As such, our efforts here are focused on providing evidence for the feasibility of STM as a general mechanism for pattern formation, rather than reproducing the specific biological details accurately.

A key guiding principle in the engineering of multicellular objectives is that cell-state trajectories should flow into attractors (i.e. stable cell-states) that reflect each cell’s appropriate cellular neighborhood along key milestones of morphogenesis (**Fig 2B**). For a demonstrative minimal system, let us consider the cell-state as fully determined by the expression levels of all its encoded ligands and receptors. The multicellular objective has access to these internal state variables, along with any signaling information conveyed by the repertoire of expressed receptors engaging extracellular ligands. Cell-state dynamics are driven by the stochastic tuning mechanism (**Fig 1**) that, over time, maximizes the multicellular objective. The objective would, thus, have multiple local maxima, each representing a single cell-state attractor (cell-types), with its corresponding optimal cellular neighborhood as perceived by its expressed receptor repertoire engaging the local ligand field. As such, the developmental trajectory of each cell corresponds to an optimization path in which the expression of ligands and receptors are driven by stochastic tuning to achieve a local maximum of the objective (**Fig 2B**). This is a local indication that the cell has achieved the appropriate expression of key cell-state variables consistent with its perceived multicellular neighborhood. As we show below, this conception leads to a simple heuristic for engineering multicellular objectives that drive the generation of desired patterns.

Given that each attractor state corresponds to one of multiple/many local maxima of the objective, the full objective can be constructed by adding multiple sub-objective terms (**equation 1**), each of which becomes exclusively dominant for a particular combination of cell-state and signaling input, and, thus, stabilizes cell state trajectory towards the corresponding attractor (cell-type). Other terms would be dominant under different cell-states and signaling inputs and, thus, drive trajectories towards other attractors. A simple strategy for generating such mutually exclusive dominant terms is to have each term be proportional to the cell-state variable(s) with the desired extreme value(s) for that cell-type, conditioned on the proper sensory input for that cell-type. For example, **equation 2** corresponds to a sub-objective (attractor) with high *intracellular* expression of ligand 1 (*L*_*1*_) in cells with high signaling input engaging receptor 2 (*R*_*2*_) by *extracellular* ligand 2 (*S*_*2*_). In our convention, *L*_*i*_ refers to the intracellular expression level of the gene encoding ligand *i* and *S*_i_ refers to the extracellular concentration of ligand *i* which is perceived by its cognate receptor *R*_*i*_.

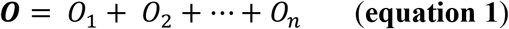

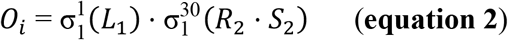

Similar to Hill functions^45^, the logistic sigmoid function (*σ*) provides a convenient way to represent biochemically realistic behaviors for the underlying molecular implementations, with saturation at high values, and parameters (*h*) reflecting steepness, and (*k*) half-maximal activity (**Fig 3A**). At low steepness (values of *h~*1, as in the first term of equation 2), the sigmoid is linearly proportional to its argument in the range of ~*k* but saturates at higher values (**Fig 3B**). The shallow steepness of this sigmoid term facilitates efficient gradient ascent optimization by stochastic tuning, leading to maximization of its argument (*L*_*1*_). As such, we refer to this first term (with *h*=1, *k*=1) as a ‘driving’ component for *L*_*1*_. On the other hand, at high values of *h*, the second sigmoid term of equation 1 (with *h=*30, *k=*1), behaves like a step function (**Fig 3B)**, implementing a binary switch at threshold *k=*1. We refer to this second sigmoid term as a ‘sensing’ component, given that its argument (*R*_*2*_ *\* S*_*2*_) senses the extracellular concentration of ligand 2 by receptor 2. We would, thus, expect that the optimization of equation 2 by stochastic tuning leads to increase in *L*_*1*_ expression only in cells that have above threshold (*k*=1) signaling from ligand 2 from neighboring cells (*R*_*2*_ *\* S*_*2*_). A key attribute of this formulation is that the higher the expression level of *L*_*1*_ in a cell, the higher the dominance of this term in the overall objective for that cell. This positive feedback naturally leads to attractor dynamics that drive cell states towards a local maximum of the overall objective.

**Fig. 3.**
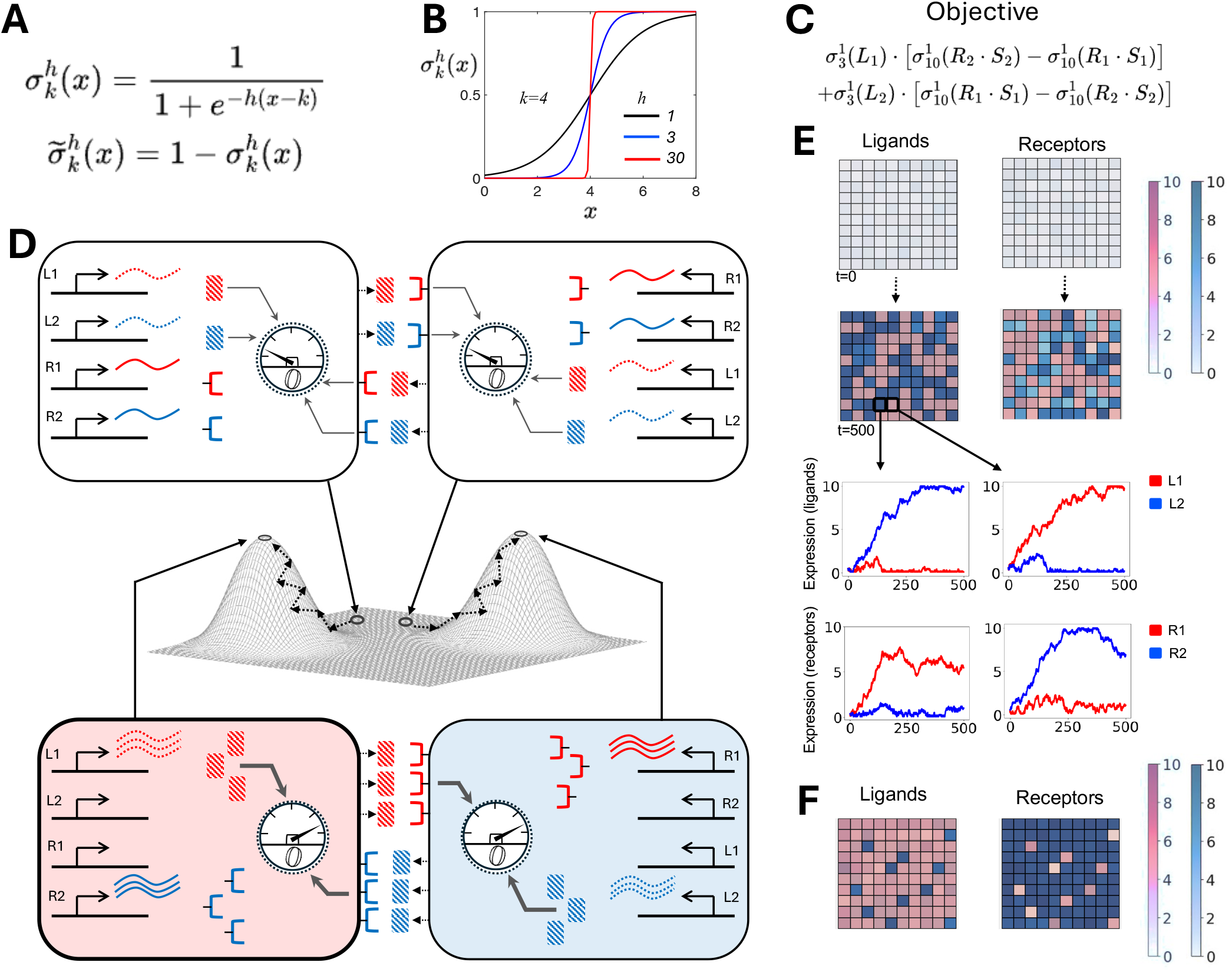
STM model of differentiation. (**A**) Logistic sigmoid function and its complement with steepness (*h*) and half-maximal activity (*k*) parameters. (**B**) behavior of the sigmoid function for different values of *h*. (**C**) Objective function for driving differentiation towards two mutually exclusive fates. (**D**) Cartoon depiction of two adjacent cells, initially expressing diffusible ligands *L*_*1*_ and *L*_*2*_ and receptors *R*_*1*_ and *R*_*2*_ at low levels. Initial small differences due to noise, drives cell states towards two distinct optima corresponding to the two optima of the objective function, here depicted as peaks on the objective landscape. (**E**) Simulation of this system for a 10×10 field of cells, with initially low expression of encoded ligands and receptors. Following 500 epochs of the simulation, cells are driven towards two mutually exclusive optima of the objective function. Expression trajectories of ligands and receptors are shown for two adjacent cells, demonstrating noisy convergence to high and low states. (**F**) Scaling the first term of the objective by 2.0 leads to imbalanced differentiation favoring the corresponding optimum.

To demonstrate the key properties of STM, let us explore a simple model of multicellular differentiation driven by the objective function of **equation 3** (**Fig 3C**):

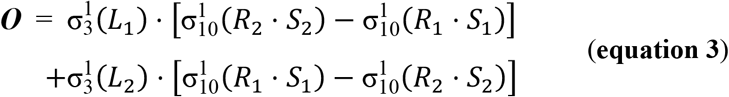

The cells of this system encode two diffusible ligands and their corresponding cognate receptors. The objective is a function of the intracellular levels of the diffusible ligands (*L*_*1*_ and *L*_*2*_) and signaling from receptors *R*_*1*_ and *R*_*2*_, engaging *extracellular* ligands 1 and 2 (*S*_*1*_ and *S*_*2*_). Optimization of this objective drives cell-state dynamics towards two distinct attractors, each corresponding to the dominance of one of the two terms, leading to mutually exclusive expression of ligands and receptors in neighboring cells. The first term of the objective is the product of two components. The first component is a sigmoid function, with a shallow steepness (*h =*1), that functions as the driving component for expression of ligand *L*_*1*_. The second component is the difference of two sigmoid functions that are sensory, first for the product of receptor *R*_*2*_ and extracellular concentration of its cognate ligand *L*_*2*_: (*R*_*2*_ * *S*_*2*_), and the second for the product of receptor *R*_*1*_ and extracellular concentration of its cognate ligand *L*_*1*_: (*R*_*1*_ * *S*_*1*_). The second term of the objective has the same functional form but with a driving component for *L*_*2*_ expression and a reversal in the order of the two sensing components. The objective of **equation 3** has a simple logic. For any cell, if the perceived signaling from ligand 2: (*R*_*2*_ * *S*_*2*_) is higher than from ligand 1: (*R*_*1*_ * *S*_*1*_), the first term dominates, driving the expression of ligand 1 to its maximum by stochastic tuning (**Fig 3D**). In the opposite scenario, if the perceived signaling from ligand 1: (*R*_*1*_ * *S*_*1*_) is higher than from ligand 2: (*R*_*2*_ * *S*_*2*_), then the second term dominates, driving the expression of ligand 2 to its maximum (**Fig 3D**). Furthermore, given each term’s proportionality to the difference in signaling from ligands 1 and 2, the expression of the receptors is also driven to mutually exclusive extremes (**Fig 3D**).

The dynamics driven by this toy objective function (**equation 3**) can be visualized for a simulation of this system for a 10 × 10 field of cells (**Fig 3E**). The cells start out with low expression of all encoded genes with normally distributed variance (μ=0.2, σ =0.1). For each cell, small initial differences in signaling from ligands 1 and 2 are amplified by stochastic tuning driven optimization, leading to mutually exclusive high expression of either ligand 1 or ligand 2, and receptor 2 and 1, respectively (**Fig 3E**). It is important to emphasize that these dynamics are driven entirely by gene expression noise reinforced by a single global scalar objective (**equation 3**).

The logic of the objective in the example above is to generate randomly interspersed small patches of two different cell types (**Fig 3E**). Since each attractor (cell-type) corresponds to the dominance of one of the two terms in the objective, we can bias the system to generate more or less of each cell-type by simply scaling each term. For example, scaling the first term of the objective in **equation 3** by a factor of 2.0 drives most of the cells towards attractor 1 fate (*L*_*1*_ high, *L*_*2*_ low), with a minority of isolated cells adopting the complementary fate (*L*_*2*_ high, *L*_*1*_ low) (**Fig 3F**).

### STM drives generation of prototypic developmental patterns

In the example above, we have shown how a simple objective can drive the differentiation of a uniform field of cells towards two mutually exclusive fates. A common scenario during development is the generation of patterns in the background of a molecular non-uniformity due to preexisting morphogen gradients. Next, we demonstrate an example of such a scenario in a toy model of segment formation (**Fig 4**). The initial molecular non-uniformity is established by a stable gradient of a morphogen that is external to the system (*E*_*1*_). Otherwise, the uniform 40 × 40 field of cells starts out with low expression of all encoded genes, with normally distributed variance (μ=0.2, *σ*=0.1). The system evolves in time through stochastic tuning of all encoded genes driven by the objective function in **Fig 4A**.

The logic of this objective is to generate a multilayered structure with two initial large domains which are subsequently differentiated at their boundary with two additional thin layers between them.

**Fig. 4.**
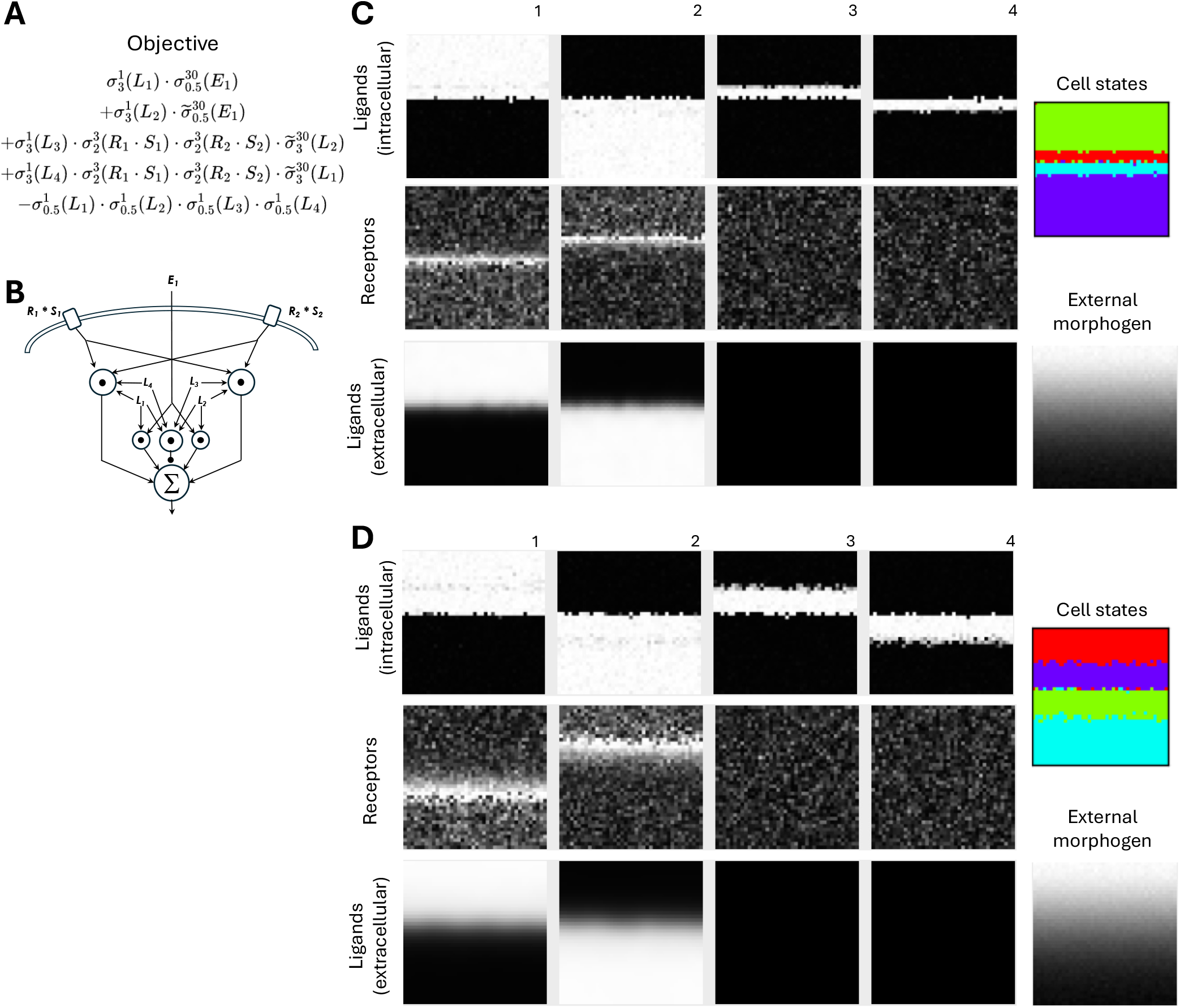
STM simulation of tissue segmentation. (**A**) Objective function. (**B**) Signaling pathway diagram corresponding to the objective. (**C**) Top: final expression of ligands 1-4 across a 40×40 field of cells. Middle: final expression of receptors 1-4. Bottom: extracellular concentrations of ligands 1-4. Only ligands 1 and 2 are diffusible. The concentration of the external morphogen gradient is also shown. Unsupervised clustering identifies distinct cell-types depicted by different colors. (**D**) Same as (C) but for a change in diffusion constants for ligands 1 and 2 from 1.0 μm^2^/s to 5.0 μm^2^/s.

As before, each term of the objective roughly corresponds to a single attractor (cell-type) which is driven by stochastic tuning to a local maximum. In addition to the external morphogen *E*_*1*_, the system has four ligands, of which only *L*_*1*_ and *L*_*2*_ are diffusible. The first term in the objective is the product of two sigmoid functions. The first sigmoid of term 1 is a driving component for ligand *L*_*1*_ since its shallow slope (*h =*1) facilitates efficient gradient ascent optimization of *L*_*1*_ expression. The second sigmoid of the first term is a sensing component and its sharp slope (*h=*30) functions to limit the dominance of the term to conditions where the extracellular morphogen *E*_*1*_ is higher than the threshold (*k*=0.5). This first term, thus, leads to increase of *L*_*1*_ expression only in cells that are exposed to a concentration of the external ligand *E*_*1*_ higher than 0.5. The second term of the objective is also a product of two sigmoids. The first sigmoid is a driving component to increase *L*_*2*_ expression and the second sigmoid complement function (**Fig 3A**) limits this activity to cells that are exposed to a concentration of external ligand *E*_*1*_ *lower* than 0.5. In combination, these first two terms drive the mutually exclusive expression of *L*_*1*_ and *L*_*2*_ across the threshold of 0.5 in the concentration of the external morphogen gradient *E*_*1*_. Ligands *L*_*1*_ and *L*_*2*_ are diffusible and, thus, establish their own extracellular concentration gradients (**Fig 4C**).

The third term of the objective is the product of four sigmoid functions. The first sigmoid is a driving component for ligand *L*_*3*_. The second and third components are sensing terms that limit the activity of the full term to the boundary between the two large *L*_*1*_/*L*_*2*_ domains, where cells perceive signaling from both diffusible ligands *L*_*1*_ and *L*_*2*_. The fourth component is a sigmoid complement function that limits the full term’s activity to the *L*_*1*_/*L*_*2*_ boundary sub-region where *L*_*2*_ expression is below 3.0. The fourth term of the objective has a similar form, but its purpose is to drive ligand *L*_*4*_ expression in cells at the same *L*_*1*_/*L*_*2*_ boundary, but limited to the sub-region where *L*_*1*_ expression is lower than 3.0. Finally, the fifth (negative) term imposes a global penalty for simultaneous expression of multiple ligands in the same cell, thus canalizing the attractor dynamics.

The form of the objective function in **Fig 4A** can be mapped to a signaling pathway diagram, akin to standard conventions (**Fig 4B**). The diagram has signaling integration nodes corresponding to products of incoming sensory and cell-state variables, and their overall integration into a single output whose directional change guides the stochastic tuning of all the encoded genes. The dynamics orchestrated by the optimization of the objective can be seen in a simulation of this system for a 40×40 field of cells, generating a hierarchically organized sequence of distinct layers, each corresponding to a unique combination of receptor and ligand expression (**Fig 4C**; **Movie 1**). Unsupervised clustering of these gene expression sates assigns each cell to one of four separate layers, with distinct colors representing each ‘cell type’ (**Fig 4C**). In addition to the form of the objective, the detailed trajectories of pattern formation can depend on the choice of system parameters such as those governing ligand diffusion and degradation rates. For example, increasing the diffusion coefficient for ligands *L*_*1*_ and *L*_*2*_ from 1.0 μm^2^/s to 5.0 μm^2^/s in the example above, leads to broader zones of expression for ligands 3 and 4 and, thus, thicker intermediate layers (**Fig 4D; Movie 2**).

A common theme in tissue organization is a pattern of repeating functional units that fulfill modular physiological roles, for example nephrons in the kidney, alveoli in the lung, or secretory glands in endocrine and exocrine organs. The objective function in **Fig 5A** generates a prototypic pattern of such functional units. The system encodes three membrane-bound (non-diffusible) ligands and their cognate receptors. The first term of this objective (**Fig 5A**) drives the expression of membrane-bound ligand *L*_*1*_ proportional to external signaling from ligand *L*_*1*_ (*R*_*1*_ *\* S*_*1*_), and in cells that perceive low external signaling from Ligand *L*_*2*_ (*R*_*2*_ *\* S*_*2*_) below the threshold of 1.0. The second term of the objective (**Fig 5A**) has the same form but for driving the expression of the membrane-bound ligand *L*_*2*_, proportional to external signaling from ligand *L*_*2*_ (*R*_*2*_ *\* S*_*2*_), and in cells that perceive low external signaling from ligand *L*_*1*_ (*R*_*1*_ *\* S*_*1*_) below the threshold of 1.0. The third term of the objective (**Fig 5A**) drives the expression of ligand *L*_*3*_ in cells that already express ligand *L*_*1*_ above the threshold of 1.0, and also perceive external signaling from ligand *L*_*2*_ (*R*_*2*_ *\* S*_*2*_) above the threshold of 1.0. The fourth negative term penalizes concurrent expression of ligands *L*_*1*_ and *L*_*2*_ in the same cells. The coefficients α and β set the relative contribution of the corresponding terms. This objective function maps to a corresponding signaling integration diagram (**Fig 5B**).

**Fig. 5.**
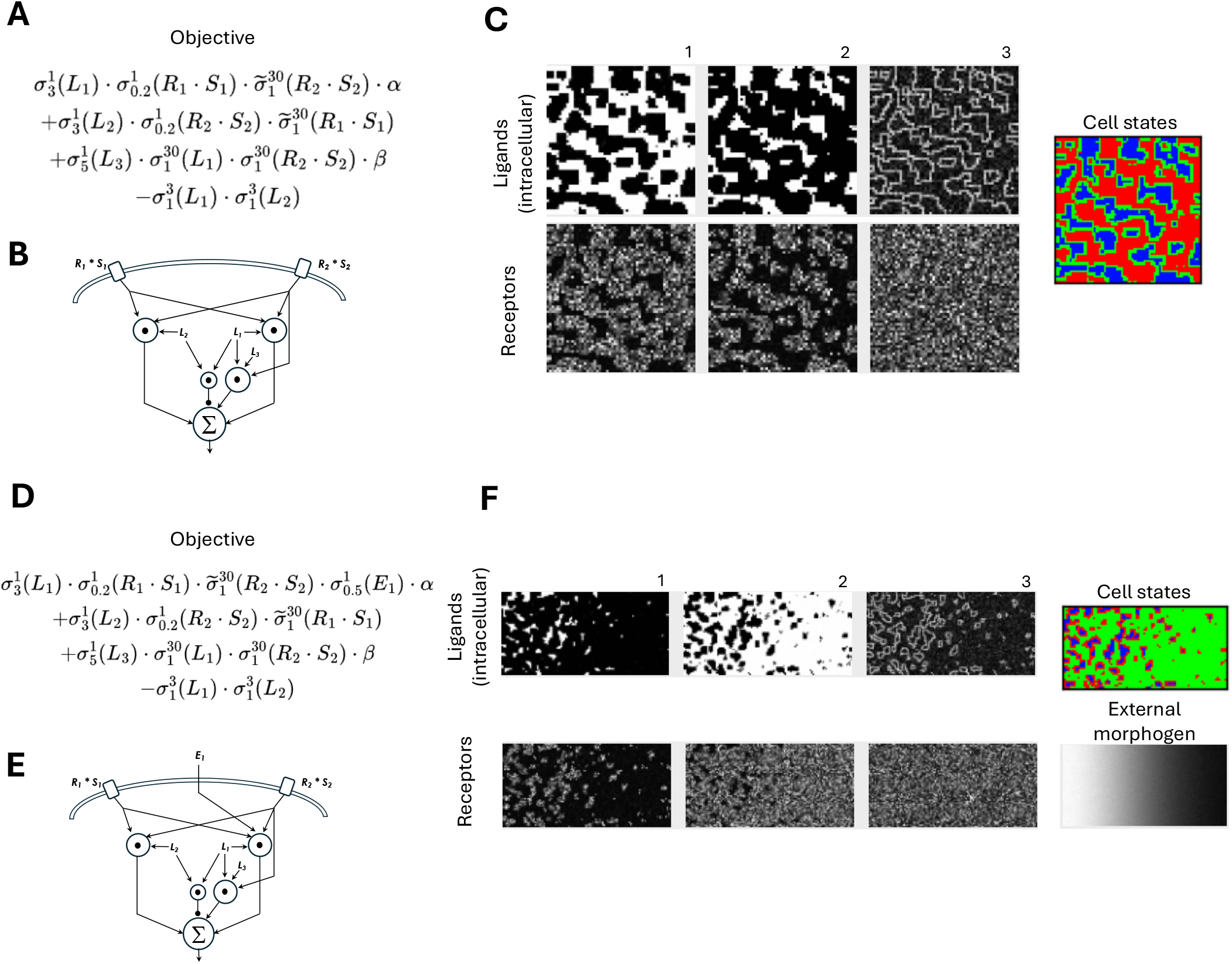
STM simulation of tissue differentiation into tubular patterns. (**A**) Objective function. (**B**) Signaling pathway diagram corresponding to the objective. (**C**) Top: final expression of ligands 1-3 across a 60×60 field of cells. Bottom: final expression of receptors 1-3. Unsupervised clustering identifies distinct cell-types depicted by distinct colors. (**D**) Objective for differentiation into tubular patterns with graded dimensions across an external morphogen gradient (*E*_*1*_). (**E**) Signaling pathway diagram corresponding to the objective in (D). (**F**) Top: final expression of ligands 1-3 across a 60 x120 field of cells. Bottom: final expression of receptors 1-3. The concentration of the external morphogen gradient is also shown. Unsupervised clustering identifies distinct cell-types depicted by different colors.

The simulation of a layer of cells driven by this objective leads to randomly distributed tubular structures emerging from a uniform background (**Fig 5C**). It is easy to see how the logic of this objective leads to the observed dynamics (**Movie 3**). The positive feedback inherent in the first two terms of the objective leads to formation of patches of cells with mutually exclusive expression of membrane-bound ligands *L*_*1*_ and *L*_*2*_ (**Fig 5C**). Subsequently, the third term drives the expression of ligand *L*_*3*_ at the boundary between the two mutually exclusive patches, in cells already expressing ligand *L*_*1*_ but also directly adjacent to cells that express the membrane-bound ligand *L*_*2*_, perceived by receptor *R*_*2*_: (*R*_*2*_ *\* S*_*2*_). This third term, thus, leads to the formation of a single-cell thickness epithelial layer between the two patches (**Fig 5C**). Coefficients α and β can be tuned to modify the size of the patches and the proper temporal order for formation of a robust epithelial layer.

The initial emergence of patches (first two terms) which are subsequently elaborated by an epithelial boundary (third term), is illustrative of the common organizational principle of hierarchical nesting in which coarse-grained patterns initially emerge at large scale and are subsequently refined through fine-grained dynamics at the smaller scale^46^. Objective-driven optimization in STM lends itself well to this scheme since a complex multi-scale process can be built-up from additive sub-objectives, each reflecting distinct phases of the process. In general, the individual terms of the objective may need to be appropriately scaled to ensure their dominance at the proper step along the hierarchically contingent process.

The modularity inherent in our formulation, where each term of the objective roughly corresponds to a single attractor state, enables easy elaboration of individual patterns and their composition into more complex forms. For example, we can add additional complexity to the basic tubular pattern (**Fig 5C**) by coupling the first term of the corresponding objective to an external morphogen gradient *E*_*1*_ (**Fig 5D,E**). As can be seen, this leads to the generation of a pattern with continuously decreasing tubular dimensions across the gradient (**Fig 5F; Movie 4**).

We can also combine multiple patterns by sharply delimiting their spatial zones of dominance using pre-existing morphogen gradients. For example, we can combine the patterns of (**Fig 3**) with that of (**Fig 5C**), by multiplying each by sharp sigmoid functions that establish mutually exclusive zones above and below the vertical morphogen *E*_*2*_ threshold of 0.50 (**Fig 6A,B**). The horizontal morphogen gradient (*E*_*1*_) acts orthogonally to modulate both patterns across the horizontal dimension, continuously decreasing tubular dimensions in the zone above *E*_*2*_=0.5, and gradually shifting the dominant interspersed cell-type below *E*_*2*_=0.5 (**Fig 6A,B**). A simulation of this system for a 120 × 80 field of cells demonstrates how such complex composite patterns can emerge from a uniform background (**Fig 6C; Movie 5**). It is important to emphasize that despite the full complexity of the objective for this system (**Fig 6A,B**), as a cell proceeds along a differentiation trajectory, progressively smaller portions of the full objective remain ‘active’ and, thus, drive the dynamics.

**Fig. 6.**
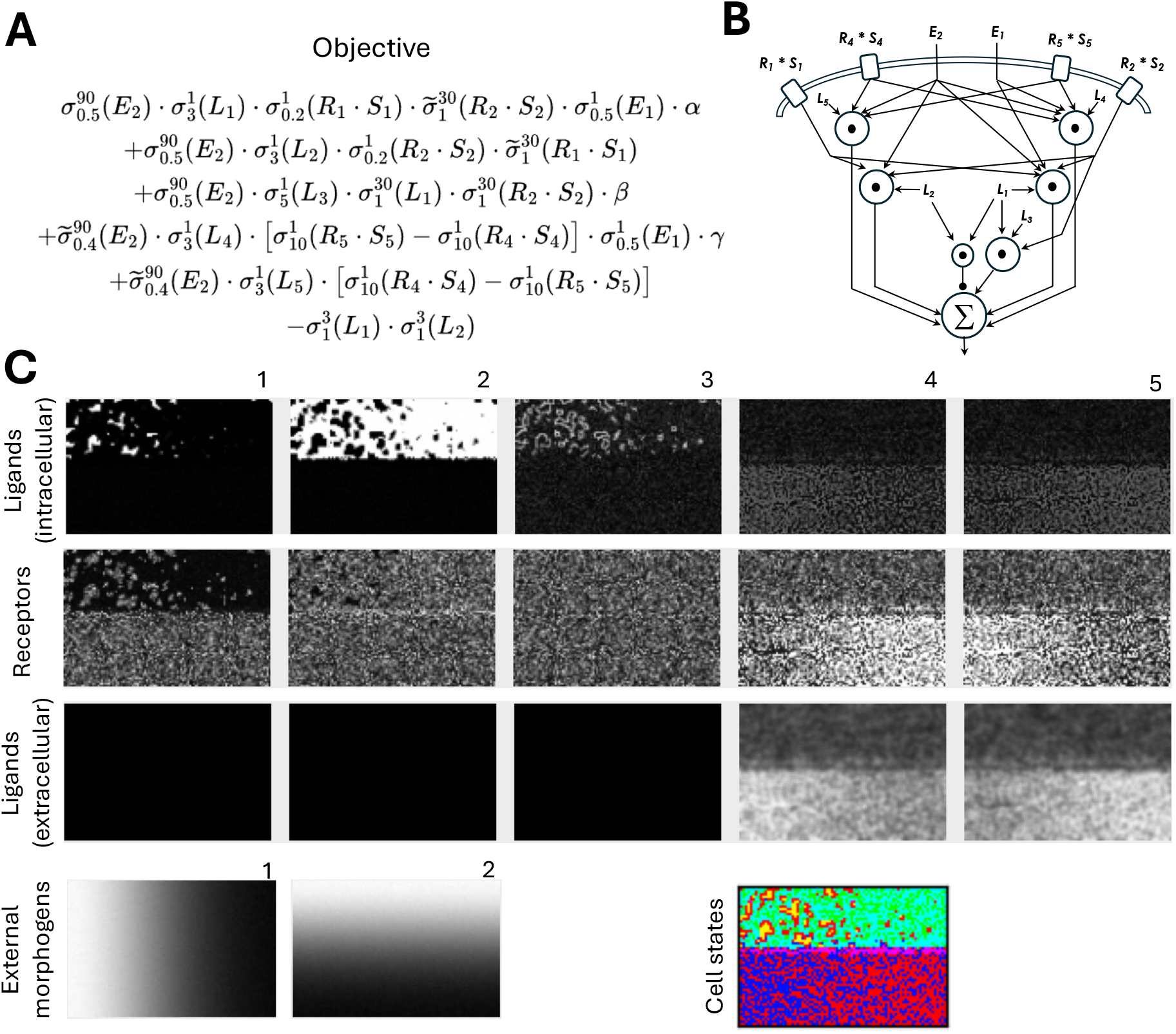
STM simulation for modular differentiation of two distinct vertical domains with horizontal variation of their features. (**A**) Objective function. (**B**) Signaling pathway diagram corresponding to the objective. (**C**) First row: final expression of ligands 1-5 across a 80 x120 field of cells. Second row: final expression of receptors 1-5. Third row: extracellular concentration of ligands 1-5. Ligands 1-3 are not diffusible. Fourth row: the concentration of two external morphogen gradients (*E*_*1*_ and *E*_*2*_). Unsupervised clustering identifies distinct cell-types depicted by different colors.

So far, we have explored how STM can drive differentiation of a uniform field of cells into a stable spatial configuration of distinct cell types. Developing tissues can also exhibit temporary periods of oscillatory behavior, for example as seen during somitogenesis^47,48^. The objective function in **Fig 7A** and its corresponding signaling pathway (**Fig 7B**) show how delayed negative feedback can lead to such oscillatory spatiotemporal dynamics. The objective function has two terms, each becoming dominant at the two spatial extremes defined by either high or low concentration of a preexisting morphogen gradient *E*_*1*_ (**Fig 7C**). The first term drives the expression of ligand *L*_*1*_ within the zone defined by *E*_*1*_ > 0.80. However, the driving term for *L*_*1*_ is also repressed by a negative term corresponding to signaling from ligand *L*_*2*_ perceived by receptor *R*_*2*_: (*R*_*2*_ *\* S*_*2*_). The constant negative term (−0.5) imposes a default drive to lower *L*_*1*_ expression. The second term of the objective drives the expression of ligand *L*_*2*_ within the opposite zone defined by *E*_*1*_ < 0.2, but only if there is significant ‘activating’ signal from ligand *L*_*1*_ perceived by receptor *R*_*1*_: (*R*_*1*_ *\* S*_*1*_). Again, the constant negative term (−0.5) imposes a default drive to lower *L*_*2*_ expression. The system starts out with low expression of all ligands and receptors. Then, *L*_*1*_ is driven to high levels in the spatial zone where *E*_*1*_ > 0.8. Over time, the diffusion of extracellular ligand *L*_*1*_ from this zone leads to the activation of the second term within the opposite zone defined by *E*_*1*_ < 0.2, increasing the expression of ligand *L*_*2*_ in these cells. However, the extracellular diffusion of ligand *L*_*2*_ from this zone back to the *L*_*1*_ high zone, leads to repression of ligand *L*_*1*_ expression. As ligands degrade over time, the *L*_*2*_ high zone is subsequently repressed since it no longer senses signaling from extracellular ligand *L*_*1*_. As can be seen (**Fig 7C**; **Movie 6**), optimization of this objective by STM leads to oscillations within the two mutually exclusive zones for expression of ligands *L*_*1*_ and *L*_*2*_.

**Fig. 7.**
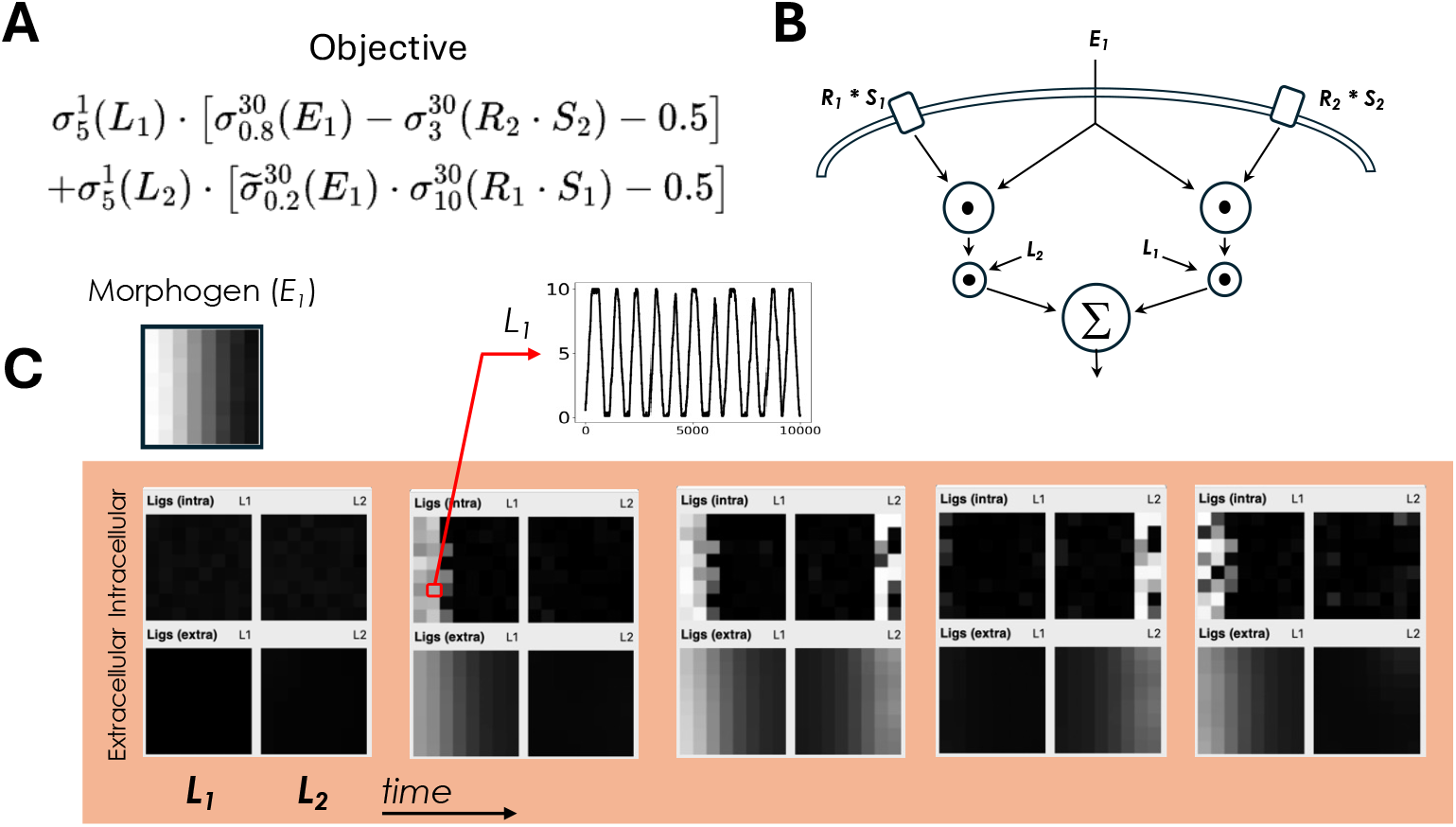
Simulation of oscillatory spatiotemporal dynamics. (**A**) Objective function for oscillatory dynamics in ligand expression at opposite extremes of the external morphogen gradient *E*_*1*_. (**B**) Signaling pathway corresponding to the objective in (A). (**C**) Top: concentration of the stable external morphogen gradient *E*_*1*_. Bottom: frames correspond to the intracellular expression and extracellular concentration of diffusible ligands *L*_*1*_ and *L*_*2*_ across an 8×8 field of cells. Cells start out with low expression of both ligands (first frame). Over time, *L*_*1*_ and *L*_*2*_ exhibit oscillations at the two extremes defined by the concentration of *E*_*1*_. Oscillations in the expression of the gene encoding *L*_*1*_ is shown for one of the cells at the left boundary.

## Discussion

We have introduced stochastic tuning-driven morphogenesis (STM), an alternative conceptualization of development in which patterns are generated by a stochastic search to maximize multicellular objectives. In contrast to the prevailing gene regulatory network paradigm^3^, the detailed trajectories of gene expression and cell states are not explicitly encoded by the genome. Rather, individual cells have agency to discover optimal trajectories through an objective-driven stochastic search, essentially corresponding to a form of single-cell reinforcement learning^49,50^. Our proposal for STM was motivated by compelling recent observations. Firstly, there is extensive evidence that cell-state transitions during differentiation and development are accompanied by periods of high gene expression noise, the functional significance of which remains highly perplexing^6–19,22^. Secondly, there is evidence that a noise-driven stochastic optimization process enables budding yeast cells to adapt to conditions that are beyond the capacity of their genetically encoded regulatory programs^28^. A phenomenon consistent with stochastic tuning is conserved in distantly related fission yeast^29^ and there is now growing evidence that stochastic tuning underlies non-mutational adaptation of mammalian cancer cells to chemotherapy^30–32^. We were, thus, compelled to consider whether a multicellular version of stochastic tuning may guide gene expression dynamics during development in a manner independent of conventional transcriptional regulatory programs. We, therefore, formulated a biologically plausible framework for STM that utilizes the stochastic tuning mechanism to perform gradient ascent optimization of multicellular objectives. We propose that these multicellular objectives map directly to the topology of commonly studied signaling pathways, and that their crosstalk and integration embodies the underlying computations performed. We have shown that a minimal implementation of STM, in 2D simulations, enables generation of a variety of prototypic developmental patterns, entirely in the absence of conventional transcriptional regulatory programs. We discuss the feasibility and broader implications of the proposed STM framework below.

### Feasibility of STM in the context of known molecular biology

As conceived here, the STM framework requires molecular machinery for: (**1**) integrating signaling events and cell state variables into effective computations that represent multicellular objectives; (**2**) broadcasting a global signal continuously reflecting changes in the multicellular objective; and (**3**) noise-driven optimization of gene expression based on both recent gene activity and the reinforcement signal broadcasted by the multicellular objective.

### Molecular computations underlying multicellular objectives

How feasible are molecular implementations of the proposed multicellular objectives? Molecular interaction networks are known to implement diverse computations^51–53^ enabling a variety of complex behaviors such as bacterial chemotaxis^54^. Recent work has revealed how simple modular computations can arise from context-dependent combinatorial utilization of signaling complexes^55,56^. Higher-order computations, on par with our proposed objectives, can emerge from arbitrary compositions of these modular components. There is also extensive evidence for signaling crosstalk and integration among multiple developmental signaling pathways^33–35,40–43^. The general mathematical form of our objectives maps directly to such signaling crosstalk and integration nodes.

### The nature of the multicellular objective signal

Here, we conceive of a single global multicellular objective, with a single scalar output, directing all morphogenetic processes. However, multiple sensory integration hubs, with distinct outputs, could also simultaneously operate to optimize cell-state trajectories, each with their own independent objective signals and batteries of morphogenetic genes. A major outstanding question is the nature of such objective signals. These signals could be conveyed by the concentration of a diffusible small molecule or abundance/modification states of signaling proteins that translocate to the nucleus.

We have conceived of these signals as being globally accessible to all genes. However, they could also be targeted to a subset of genes whose functions have evolved specialized roles in morphogenesis. Such targeting could be achieved through protein-protein interactions with specialized transcriptional complexes such as the mediator^57,58^ or co-activator complexes^59,60^ that are known to have gene targeting biases. Alternatively, the signaling proteins could be targeted to specific genes through direct (sequence-specific) protein-DNA interactions. In such a scenario, it is important to distinguish the function of such objective-signal conveying DNA-binding proteins from conventional transcription factors whose DNA binding leads to deterministic activation or repression of transcription. In this context, conventional DNA enhancers^5^ could function as the convergence points of multiple objective signals, which through the combinatorial configuration of TF binding sites, may encode additional complexity of the composite objective in a locus-specific fashion.

Given the scalar nature of the objective, it could also be conveyed by changes in ionic concentrations (e.g. Ca^2+^) or even membrane potential. Intriguingly, there is evidence that membrane potential correlates with morphogenetic transitions, and perturbations to membrane potential can affect morphogenic fates^61–63^. In fact, Levin and co-workers have suggested that bioelectric patterns may encode anatomical setpoints that organize morphogenesis during development and regeneration^64^. Whatever the nature of these signals, it is important to stress that it is the direction and scale of the recent *change* in the objective, rather than its absolute value, which is critical for gradient ascent optimization of multicellular objectives by stochastic tuning.

### Noise-driven optimization of gene expression

Cell-state dynamics in STM are driven by stochastic tuning, the phenomenon in which gene expression is driven towards optima by the reinforcement of random transcriptional changes^28^ (**Fig1**). This molecular form of gradient-ascent^44^ requires: (**1**) transcriptional noise; (**2**) local memory of recent transcriptional activity; and (**3**) modulation of transcriptional output based on both recent transcriptional activity *and* the direction of currently observed change in the multicellular objective. How feasible are these ingredients? Random transcriptional bursting is a ubiquitous feature of eukaryotic gene expression, especially in developmental genes where its role remains puzzling^13–16^. Transcriptional memory is also a given, as co-transcriptional histone modification can locally store recent memory of transcriptional activity^65,66^. Given that histone modifications can, in turn, affect the binding and function of chromatin factors, and consequent transcription^67–69^, the machinery for context-dependent reinforcement mechanism is clearly feasible. Indeed, the evolution of elaborate multivalent chromatin modification states, dynamically altered by dozens of effectors, suggests that chromatin implements complex algorithms beyond our current appreciation^70^. It is notable that mutations to chromatin-related regulators manifest disproportionately in developmental phenotypes^71–74^. Irrespective of detailed molecular mechanisms, there is now growing evidence for stochastic tuning in the adaptation of distantly related eukaryotes, including mammalian cells^28–32^. This existing stochastic tuning mechanism can, thus, be coopted to drive developmental gene expression by coupling to multicellular objectives, as we have demonstrated here in our simulations.

### How do we reconcile STM in the context of the dominant gene regulatory network paradigm?

Since the pioneering work of Jacob and Monod, more than sixty years ago, cellular behaviors are thought to rely on predefined regulatory programs established by dedicated genetically encoded pathways^75^. The Jacob and Monod framework is such a foundational organizing principle that we rarely consider its wide-reaching influence^76^. STM represents a fundamental departure from Jacob and Monod. Indeed, the proposition that individual cells can, instead, utilize a noise-driven trial and error process to *empirically* establish cell states goes against our most cherished notions of cellular function and behavior. Is there a way to reconcile these highly divergent conceptions? We briefly discuss some possibilities below.

#### GRNs and STM function in distinct specialized roles

STM and conventional GRNs may have evolved complementary, specialized, roles during distinct phases of development. For example, GRNs may drive early morphogenesis, establishing coarse-grained patterns, and STM would later dominate in the formation of small-scale features. The two mechanisms may also be contrasted in terms of timescale. The deterministic nature of GRNs make them better suited for dynamics on a rapid timescale, whereas the search process inherent in STM is expected to unfold with a slower cadence. As such, organisms that have evolved fast developmental life histories may disproportionately depend on GRNs relative to STM. It is notable that, for convenience, molecular studies of development have been largely focused on such rapidly developing species, including *Drosophila melanogaster* and *Caenorhabditis elegans*, and our mechanistic inferences may, thus, reflect this bias.

#### GRNs and STM work in concert

GRNs and STM may be concurrently operative at all stages of development. For example, GRNs may establish initially imprecise patterns of gene expression, and STM functions to fine-tune them. Alternatively, STM may be the ancestral mechanism and GRNs may have later evolved to canalize the dynamics, enhancing both the speed and robustness of developmental trajectories. In such a scenario, experimental studies may reveal strong correlations between transcription factor-DNA interactions and developmental gene expression. However, these interactions, by themselves, may be auxiliary and not the *key* causal driving events, even though their experimental perturbations may, nevertheless, partly contribute to the observed changes in gene expression.

#### STM is the primary underlying mechanism

A tantalizing possibility is that STM is the primary underlying mechanism of developmental gene expression, and that the deterministic GRN abstraction is our best approximation of an incompletely understood process. If this is true, how do we reconcile the vast evidence for the role of dedicated transcription factors in regulating developmental gene expression^5^? As discussed above, the multicellular objective signals may be conveyed in a targeted manner by transcription factor-DNA interactions at promoters/enhancers of genes with roles in morphogenesis. However, in reality, and contrary to the widely accepted GRN paradigm, these interactions do not *deterministically* activate/repress gene expression. Rather, these transcription factors convey valences reflecting changes in optimality of the multicellular objective, and the actual direction and scale of change in gene expression depends on both the conveyed valence and the recent history of transcriptional change at the locus. In this way, there is no consistent causal effect, up or down, of a specific TF-DNA binding event on gene expression, even though, in an isolated context, observations may, nevertheless, show a strong correlation one way or another. As such, the STM mechanism may have remained hidden to experimental studies due to our strong epistemological bias to infer a *deterministic* process consistent with the Jacob and Monod framework.

Recent observations that differentiation and development are accompanied by significant gene expression noise^6–19^ challenge the deterministic Jacob and Monod paradigm. Indeed, often the relationship between signaling pathway activation, TF nuclear localization, and target gene expression is not characterized by a simple deterministic cascade of activating or repressing events. In fact, careful studies of these cascades are revealing complex relationships between signaling pathway activation and gene expression, often mediated by temporally complex non-deterministic dynamics^77–86^ and TF nucleo-cytoplasmic oscillations^77,87–93^ of perplexing functional utility. In the context of STM, these observations may be more consistent with a mechanism for noise-driven, objective-reinforced, optimization of gene expression, rather than a precise deterministic cascade of genetically encoded regulatory events.

### STM disruption in etiology of disease

Here, we have focused on the potential role of STM in differentiation and development. However, STM may also operate in mature organs, maintaining proper tissue organization in the face of mechanical or physiological perturbations. As such, STM may be a key factor in disease pathogenesis. There is now growing evidence for the role of stochastic tuning-based adaptation in the ‘plasticity’ of tumor cells responding to chemotherapy^30–32^. It is intriguing to consider whether stochastic tuning-driven morphogenesis may similarly play a role in the process of cancer initiation, as disruption of local signaling events, for example in the setting of inflammation, may push cell-states beyond the capacity of STM to stabilize. Indeed, increasing evidence for lack of common driver mutations in many cancers^94–97^, and corresponding observations of ‘driver’ mutations in pathologically normal tissues^98^, are consistent with cancer initiation by a non-mutational process^99^. As such, the disruption of STM-based multicellular homeostasis, due to pathological processes such as inflammation, may be a key contributor to cancer initiation. In such a scenario, nominally driver mutations do not play the central ‘driving’ role. Such mutations could either modify the multicellular objective landscape, making destabilized neoplastic trajectories more probable and/or accumulate, early on, *following* STM epigenetic destabilization, as tumor cells evolve higher reproductive fitness and immune evasive potential.

### Future directions

Noise is often considered an irreducible nuisance in biological processes. In the context of STM, noise becomes the essential driving force for an empirical search to maximize multicellular objectives. We have shown that a minimal system based on STM can generate prototypic patterns reminiscent of differentiating tissues. Whether, and to what extent, real morphogenesis relies on STM must be explored in future experimental work. We need to extend the evidence for non-deterministic behaviors in model developmental systems and to see whether observations of signaling and developmental gene expression are more consistent with a stochastic tuning based optimization process. Recent work on combinatorial logic of developmental signaling pathways^55,56,100^ has revealed computations that map directly to component inputs of our proposed multicellular objectives. How these component inputs converge to signaling integration hubs, the effective functions computed by these hubs, the nature of the emerging multicellular objective signals, and whether they function to reinforce noise-driven developmental gene expression are high priorities to explore in future work.

Here, we developed the simplest possible system for simulating the essential features of STM. As such, we restricted cell-state representations to ligands and receptors, as they are the key drivers of multicellular objectives. However, the expression of these factors can, in turn, activate downstream batteries of effector genes that establish distinct cell biological features specific to each cell type. We have developed a simple heuristic for writing objective functions that generate a set of exemplary differentiation patterns. Nevertheless, the general mapping between these objectives, the attractor landscapes they create, and the tissue patterns they lead to, is a rich topic for future exploration. Furthermore, we have focused on morphogenesis of molecular patterns in static two-dimensional arrays of cells. However, the STM framework can be adapted to drive mechanical morphogenesis^101^ in more realistic three-dimensional cell assemblies. This will require elaboration of our simulations with additional features, including proliferation, polarization, mechanosensory transduction, and cell movements. This is a promising direction for future work.

## Acknowledgements

We thank the Tavazoie laboratory for helpful discussions. S.T. is supported by a grant from NIH/NIGMS (R01GM139215). JT was supported by the Amgen Scholars program for summer research at Columbia University.

## Author Contributions

S.T. conceived the study and developed the theoretical framework. J.T. and S.T. designed algorithms and developed code. S.T. performed analyses and generated figures. S.T. wrote the paper. J.T. and S.T. reviewed and edited the paper.

## Methods

### Numerical simulation and graphical display of stochastic tuning-driven morphogenesis

Numerical simulations were performed by implementing the STM model into an interactive graphical user interface (GUI) using the Python programming language^102^. The Tkinter-based interface^102^ allows for fast setup and exploration of STM in 2D lattices of interacting cells (see **Figure M1** below) Objective functions, parameters, and constraints can be easily specified directly in the interface or saved and then loaded from YAML files. To simulate large systems, the interface and underlying simulation makes use of NumPy^103^ and Numba^104^ for just-in-time (JIT) compiling and vectorization of array operations. All simulations were performed using Python 3.8. Access to the GUI with example simulations is provided in the source code at: Zenodo: **10.5281/zenodo.15698193**

**Figure M1:**
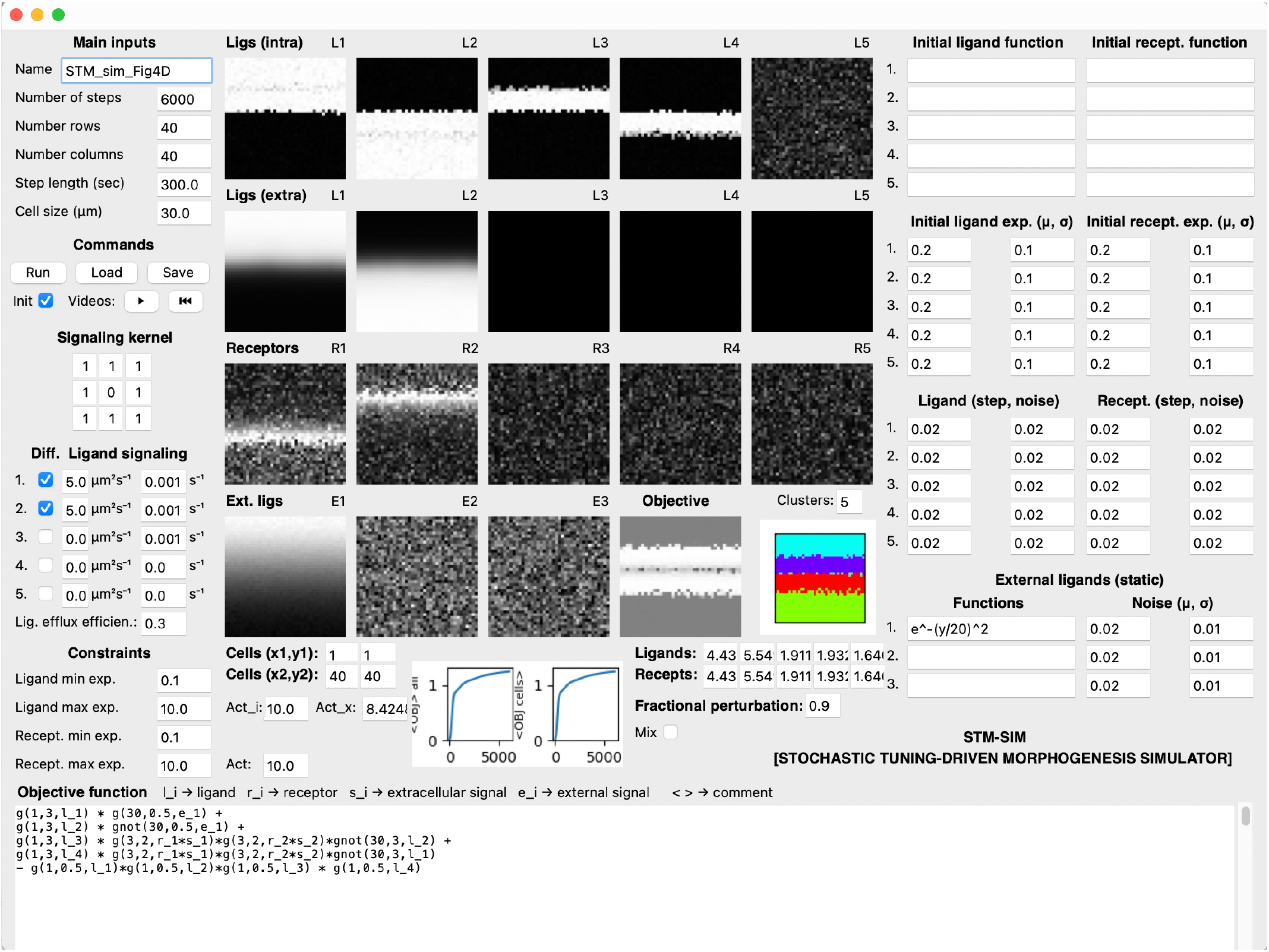
GUI for Simulation of 2D stochastic tuning-driven morphogenesis.

#### To setup the environment with Anaconda

$ conda create --name STM_*simulation* python=3.12

$ conda activate STM_*simulation*

#### To run the simulation GUI

$ pip install -r requirements.txt

$ python main.py

### Simulation GUI input parameters

#### Main inputs

The name of the simulation provides an identifier for saving/loading GUI configurations described below. High-level simulation specifications such as the size of the environment, number of simulation steps, simulation step-length, and cell-size parameters are entered here. Unless stated otherwise, simulation step length (epoch) was 300 seconds and cell-size was 30 μm.

#### Commands

This section provides GUI controls to run an instance of the STM model, load/save the GUI configuration, and play/rewind the graphics visualizing the dynamics of pattern formation in terms of the intracellular expression of ligands, receptors, and the extracellular concentration of diffusible ligands and external morphogens. The GUI also outputs the matrix of the final objective values for each cell, and the average objective of all the cells combined, along the simulation steps. We have also provided the ability to designate a rectangular section for a subset of cells (with beginning and ending x,y coordinates), inside of which the average objective is also plotted. In addition, the average values of all the ligands and receptors in this designated region is displayed, along with the capacity to perturb this sub-region, by either mixing all the values randomly, or performing a random fractional (0-1) perturbation on their final values and re-running the simulation from the previous endpoint by unclicking the ‘Init’ flag from within the Command section.

#### Signaling kernel

A 3×3 neighborhood (eight neighbors and self) can be specified to control how nearby fixed (membrane-bound) ligands interact with each cell.

#### Diffusible ligand signaling

Ligands are indicated diffusible (or fixed to the cell membrane) by checking the respective ligand number, providing diffusion coefficients, and specifying the degradation rate. Additionally, a global ligand efflux efficiency parameter (0-1) determines the efficiency with which intracellular ligands are excreted. For example, in simulation of Figure 4C, ligands 1 and 2 are diffusible with diffusion coefficients of 1.0 μm^2^/s and degradation rates of 0.001 s^-1^. The ligand efflux efficiency parameter was set at 0.3.

#### Constraints

Nondimensional constraints on expression bounds for all ligand/receptors can be applied in this section. For example, in simulation of Figure 4, Ligand_min_ = 0.1; Ligand_max_ = 10; Receptor_min_ = 0.1; Receptor_max_ = 10.

#### Objective function

Custom functions describing how ligands, receptors, and signals contribute to each cell’s objective are provided here. For ease of specifying objectives, logistic functions can be used, corresponding to the sigmoid: *g(h,k,x)*, and sigmoid complement (1-sigmoid): *gnot(h,k,x)* functions (see **Fig 3A,B**). The intracellular expression level of the gene encoding ligand *i* is represented by l_i; the extracellular concentration of ligand *i* is represented by s_i; and the expression level of receptor *i* is represented by r_i. For example, for the objective function of Figure 4, the objective function is expressed as:

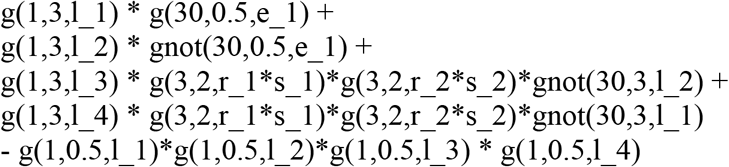

#### Initial ligand/receptor functions

Here, functions can be entered to describe the initial conditions for intracellular ligand/receptor expression with added Gaussian noise.

#### Ligand/receptor tuning

The nondimensional discrete step size and noise for the expression modulation of each ligand and receptor by stochastic tuning are parameterized in this section. For all simulations, the scale of step-size and noise were set equal. For simulation of Figure 4, step amplitude = 0.02; noise amplitude = 0.02.

#### External ligands

Static external morphogen gradients simulated from cells outside the environment are specified by functions evaluated at each cell’s position (in μm) in the environment with added noise, if desired. For example, for the simulation of Figure 4, the following stable external morphogen gradient was used with added noise (μ=0.02; σ=0.01):

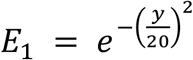

#### Unsupervised cell-type identification

To label the emergent cell types, following the completion of a simulation, unsupervised (*k-*means) clustering is performed on the expression of all the genes using *cluster*.*KMeans* from Scikit-learn^105^. Within the STM interface, one can specify the number of desired clusters (expected cell types) to be identified, which are represented by distinct colors. For robust identification of dominant cell-types, users should specify the ‘Clusters’ parameter to be one or two higher than the expected number of distinct cell-types.

#### Stochastic tuning of gene expression

For each step within the simulation, each cell will stochastically tune the transcriptional output of each gene *E*_*i*_ based on the following equation, with step-amplitude (*k*_i_) and discrete random noise amplitude (+/-|*η*_*i*_|) parameters (see above). For simplicity, all genes have the same mRNA degradation and translation rates, making protein products proportional to *E*_*i*_. Also see Freddolino et al.^28^ and **Figure 1** for more details on the stochastic tuning model.

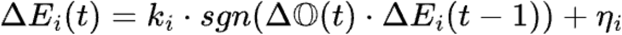

#### Ligand diffusion

Each cell is defined as a 2D square with side length *Δl* (parameter cell-size). For a lattice of *N × M* cells, we define the environment *E* = [*0, NΔl*] × [*0, MΔl*]. Concentrations (*u*_*j*_) of individual ligands *j* are computed with the following partial differential equation, with diffusion coefficient (*α*_*j*_) and incorporating the

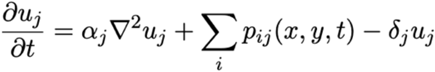

contribution of diffusible ligands *p*_*ij*_ from each cell *i* and ligand degradation rates *(δ* _*j*_).

Concentrations are solved numerically with a 2D Forward Time-Centered Space scheme with Neumann boundary conditions:

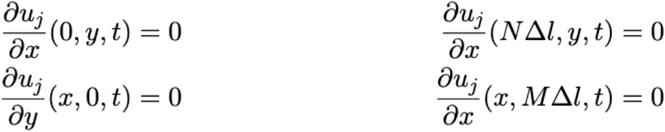

For fixed membrane-bound ligands, we convolve nearby cells with a default kernel of *K*, which can be further defined within the interface.

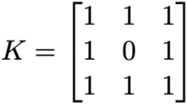

#### Parameters for simulations

For all simulations, cell-size was set at 30 μm and epoch duration (step-length) was set at 300 seconds. Other parameter choices are shown below.

### Fig 3

#### Ligand and receptor bounds

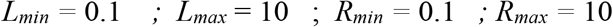

#### Ligand diffusion and degradation

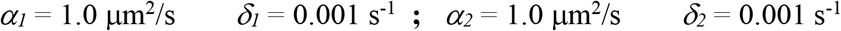

#### Stochastic tuning parameters (step amplitude and noise amplitude)

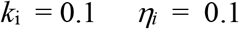

### Fig 4

#### Ligand and receptor bounds

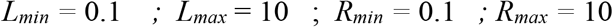

#### Ligand diffusion and degradation

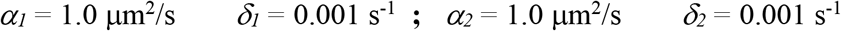

#### Stochastic tuning parameters (step amplitude and noise amplitude)

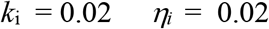

### Fig 5

#### Ligand and receptor bounds

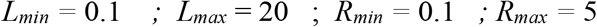

#### Ligand diffusion and degradation

No diffusible ligands

#### Stochastic tuning parameters (step amplitude and noise amplitude)

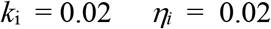

### Fig 6

#### Ligand and receptor bounds

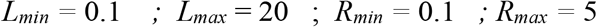

#### Ligand diffusion and degradation

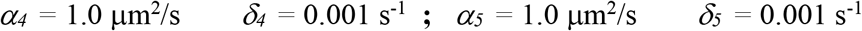

#### Stochastic tuning parameters (step amplitude and noise amplitude)

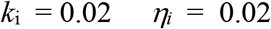

### Fig 7

#### Ligand and receptor bounds

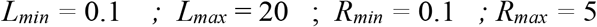

#### Ligand diffusion and degradation

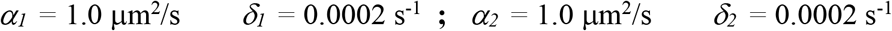

#### Stochastic tuning parameters (step amplitude and noise amplitude)

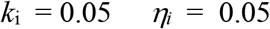

